# Membrane asymmetry facilitates murine norovirus entry and persistent enteric infection

**DOI:** 10.1101/2024.11.06.622376

**Authors:** Brittany M. Stewart, Linley R. Pierce, Mikayla C. Olson, Robert C. Orchard

## Abstract

Norovirus, the leading cause of gastroenteritis worldwide, is a non-enveloped virus whose tropism is determined in part by the expression patterns of entry receptors. However, the contribution of cellular lipids to viral entry is not well understood. Here, we determined that the asymmetrical distribution of lipids within membrane bilayers is required for murine norovirus (MNV) replication. Specifically, TMEM30a, an essential subunit of lipid flippases, is required for MNV replication in vitro. Disruption of TMEM30a in mouse intestinal epithelial cells prevents persistent, enteric infection by MNV in vivo. Mechanistically, TMEM30a facilitates MNV binding and entry. Surprisingly, exoplasmic phosphatidylserine (PS), a typical marker of dying cells, does not inhibit MNV infection. Rather, TMEM30a maintains a lipid ordered state that impacts membrane fluidity that is necessary for the low affinity, high avidity binding of MNV to cells. Our data provides a new role for lipid asymmetry in promoting non-enveloped virus infection *in vitro* and norovirus persistence in vivo.

## Introduction

The plasma membrane provides an intrinsic barrier blocking viral access to the host cytosol. Research efforts have largely focused on how receptors enable viruses to cross membranes. In contrast, very little is known about how lipids regulate viral entry. Biological membranes consist of lipid bilayers with asymmetric distribution [1]. Amine-containing phospholipids such as phosphatidylethanolamine (PE) and phosphatidylserine (PS) are found on the inner leaflet of the plasma membrane, but are nearly absent on the outer leaflet. Lipid asymmetry is established and maintained by lipid flippases and floppases. Flippases transport lipids from the exoplasmic side of the membrane to the cytoplasmic side, while floppases catalyze cytoplasmic to exoplasmic transport [2, 3]. It is appreciated that enveloped viruses promote the incorporation of exoplasmic PS in viral envelopes to promote uptake by scavenger receptors [4-6]. This process, coined viral apoptotic mimicry, has been implicated for numerous enveloped viruses including Filoviruses, Orthopoxviruses, and Flaviviruses [5]. However, whether lipid asymmetry is important for non-enveloped virus entry is unknown.

Noroviruses (NoV) are non-enveloped, positive-sense RNA viruses and are a leading cause of gastroenteritis worldwide [7, 8]. Norovirus shedding has been detected in asymptomatic individuals months after symptom resolution [9, 10]. Norovirus persistence has been postulated to be the initiating source of outbreaks, yet our ability to treat or clear persistent norovirus infections is limited [11]. Murine norovirus has emerged as the preeminent HNoV model system as it is a natural pathogen of mice, replicates robustly in standard cell lines, and certain MNV strains cause persistent enteric infection of mice that models the long-term shedding of HNoV infections [12-14]. We and others have conducted genome-wide CRISPR screens to identify host genes required for MNV replication [15-17]. In addition to identifying CD300lf as a proteinaceous receptor for MNV, numerous other genes were associated with norovirus replication [15, 17]. One such gene, TMEM30a, also called CDC50a, is an essential component of P4-ATPase flippase complexes [18]. Without TMEM30a, lipid flippases are unable to leave the endoplasmic reticulum and cells lose lipid asymmetry; yet how TMEM30a promotes norovirus replication is unclear [19, 20].

Here we set out to determine how lipid asymmetry maintained by lipid flippases facilitates MNV replication. We determined that TMEM30a is required for replication in vitro and loss of TMEM30a in intestinal epithelial cells prevents persistent, enteric infection by MNV. Interestingly, despite lipid asymmetry being disrupted throughout the cell, MNV only requires TMEM30a for viral entry. In the absence of TMEM30a, MNV is unable to efficiently bind cells even when CD300lf is present on the cell surface. We present a model in which membrane order and fluidity independent of lipid head groups such as PS are important for promoting norovirus infection and persistence.

## Results

### TMEM30a is required for efficient MNV infection *in vitro* and *in vivo*

To verify that TMEM30a is a pro-viral factor for MNV, we generated TMEM30a knockout BV2 cells (BV2ΔTMEM30a) and confirmed that these BV2ΔTMEM30a cells have the anticipated increase in exoplasmic PS and PE via staining with Annexin V and Duramycin respectively (**Figure 1A and 1B**) [18, 21]. MNV strains exhibit diverse cellular and tissue tropisms in vivo [22-24]. We first tested the ability of two representative MNV strains to replicate in BV2ΔTMEM30a cells. MNV^CR6^ causes persistent infection of intestinal tuft cells while MNV^CW3^ infects immune cells causing an acute systemic infection [22, 23, 25]. Compared to wild-type BV2 cells, BV2ΔTMEM30a cells were deficient in MNV production at a single cycle of replication (12 hours) or at multiple cycles (24 hours) (**Figure 1C and 1D**). Similar results were observed with both MNV^CR6^ and MNV^CW3^. We next extended our analysis to a panel of diverse MNV strains that represent major branches on an MNV phylogenetic tree [12]. 12 hours post-infection BV2ΔTMEM30a cells produced significantly less viral titers compared to wild-type counterparts for MNV strains CW3, CR6, CR3, CR7, and S99 (**Figure 1E**). We observed a reduction, although not statistically significant, in viral titers for MNV strain S99 in BV2ΔTMEM30a cells (**Figure 1E**). Importantly, the defect in viral production for MNV^CR6^ was complemented via expression of a *TMEM30a* cDNA (**Figure 1F and 1G**). These data indicate that TMEM30a is required for optimal in vitro infection by diverse MNV strains.

**Figure 1:**
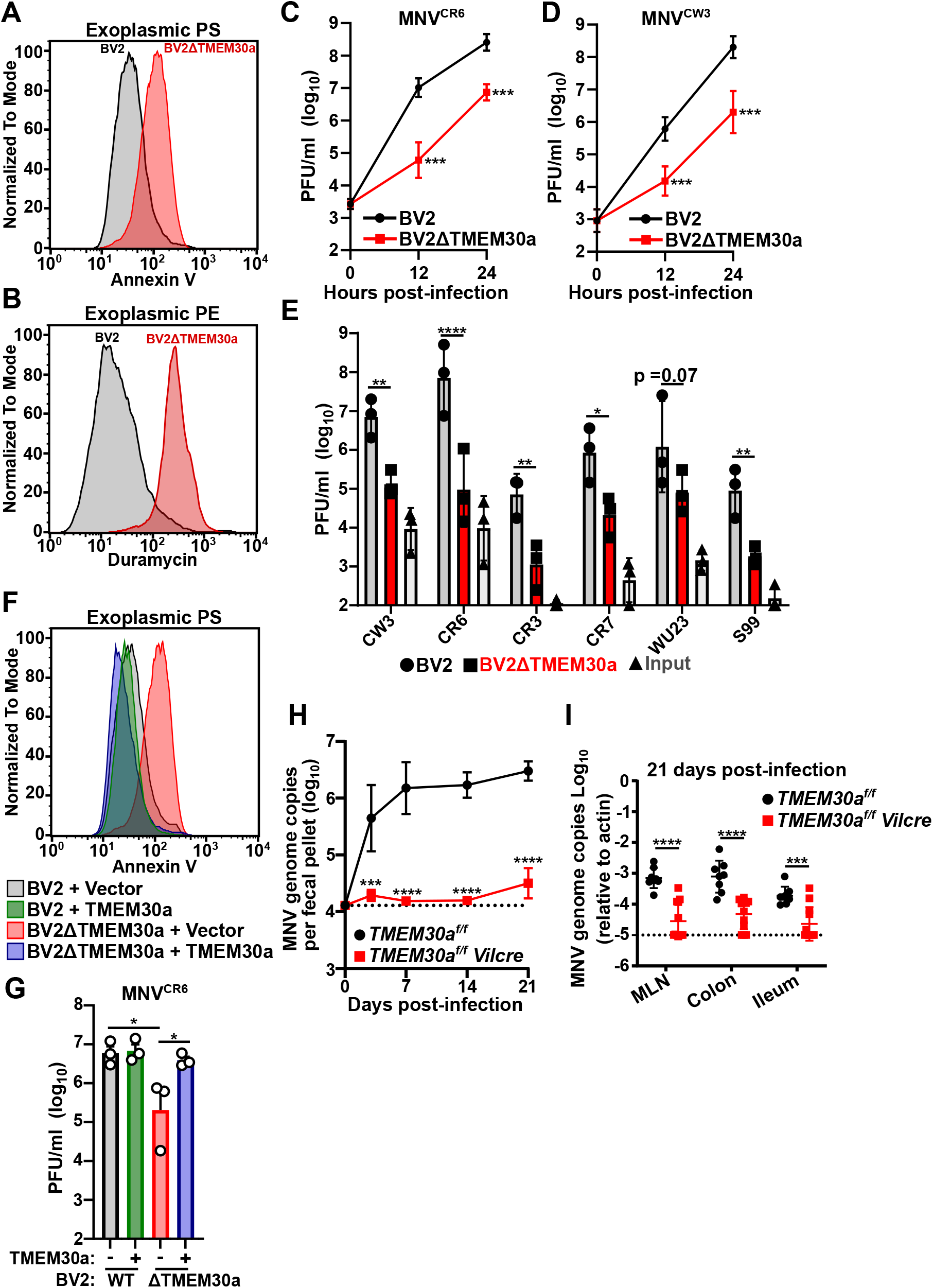
TMEM30a is required for MNV replication in vitro and in vivo. **A-B)** Representative Histograms comparing Annexin V (**A**; top) or Duramycin (**B**; bottom) staining of BV2 WT and BV2ΔTMEM30a cells. **C-D)** BV2 or BV2ΔTMEM30a cells were challenged with MNV strains MNV^CR6^ (**C**; left) or MNV^CW3^ (**D** ;right) at a multiplicity of infection (MOI) of 0.05. Viral production was enumerated using plaque assays (PFU; plaque forming units) at the indicated time points. **E)** BV2 or BV2ΔTMEM30a cells were challenged with indicated MNV strains and 12 hours post-infection viral production was enumerated using plaque assay. Viral input is also graphed. **F)** Representative Histograms comparing Annexin V staining of indicated cell lines **G)** BV2 or BV2ΔTMEM30a cells expressing either an empty vector or TMEM30a were challenged with MNV^CR6^ at an MOI of 0.05. Viral production was measured at 12 hours post-infection via plaque assay. **H-I)** *TMEMfl/fl* or *TMEMfl/fl Vilcre* mice were challenged with 10^6^ PFU *per oral* of MNV^CR6^ and genome copies were enumerated in the feces at the indicated times **(H)** or in the indicated tissues 21 days post-infection **(I)**. Each dot represents a single mouse and the dashed lines indicate the limit of detection. All data are shown as mean ± S.D. from three independent experiments and analyzed by one-way ANOVA with Tukey’s multiple comparison test. Statistical significance is annotated as follows: ns not significant, * P < 0.05, ** P< 0.01, *** P< 0.001, **** P<0.0001

Germline deletion of TMEM30a is embryonic lethal [26]. To test the contribution of TMEM30a to MNV infection in vivo, we generated conditional knockout mice (*TMEM*^*fl/fl*^) crossed to Villin Cre (*Vilcre*) to specifically target intestinal epithelial cells, including tuft cells, the reservoir for persistent MNV infections [23, 27]. Compared to cre-negative littermate controls, *TMEM30a*^*fl/fl*^*-Vilcre* mice were unable to support persistent fecal shedding of MNV^CR6^ (**Figure 1H**). Further examination of the tissues demonstrated a dramatic reduction in viral genomes in the colon, ileum, and mesenteric lymph nodes (**Figure 1I**). These data indicate that TMEM30a is required for MNV to establish a persistent infection in both tissues and fecal shedding.

### TMEM30a is required for Viral Entry and Binding Despite CD300lf at the Cell Surface

TMEM30a deficiency impacts lipid asymmetry throughout the cell and has the potential to impact multiple stages of the MNV replication cycle [28]. MNV engages cellular membranes at several points during its replication cycle including the binding of CD300lf at the plasma membrane, establishment of a replication complex via reorganization of the endomembrane system, and viral release from infected cells [29]. Equivalent levels of infectious particles were produced from wild-type BV2 and BV2ΔTMEM30a cells transfected with MNV genomes, indicating that only MNV entry requires TMEM30a (**Figure 2A**). We then tested the ability of MNV^CR6^ to bind BV2ΔTMEM30a cells. While MNV^CR6^ robustly bound wild-type BV2 cells, it was significantly impaired in binding BV2ΔTMEM30a cells or the viral receptor deficient BV2ΔCD300lf cells (**Figure 2B**). Importantly, this binding defect was rescued by expression of TMEM30a cDNA (**Figure 2B**). Because CD300lf expression is essential for MNV binding and entry, we tested whether overexpression of CD300lf could overcome MNV dependence on TMEM30a for infection. However, there was no improvement in infectious particle production in BV2ΔTMEM30a cells when CD300lf was overexpressed despite the receptor being robustly detected on the surface of the cells (**Figure 2C and 2D**). Taken together these findings suggest a model in which TMEM30a is required for the entry of MNV while all other steps of viral replication are not significantly affected.

**Figure 2:**
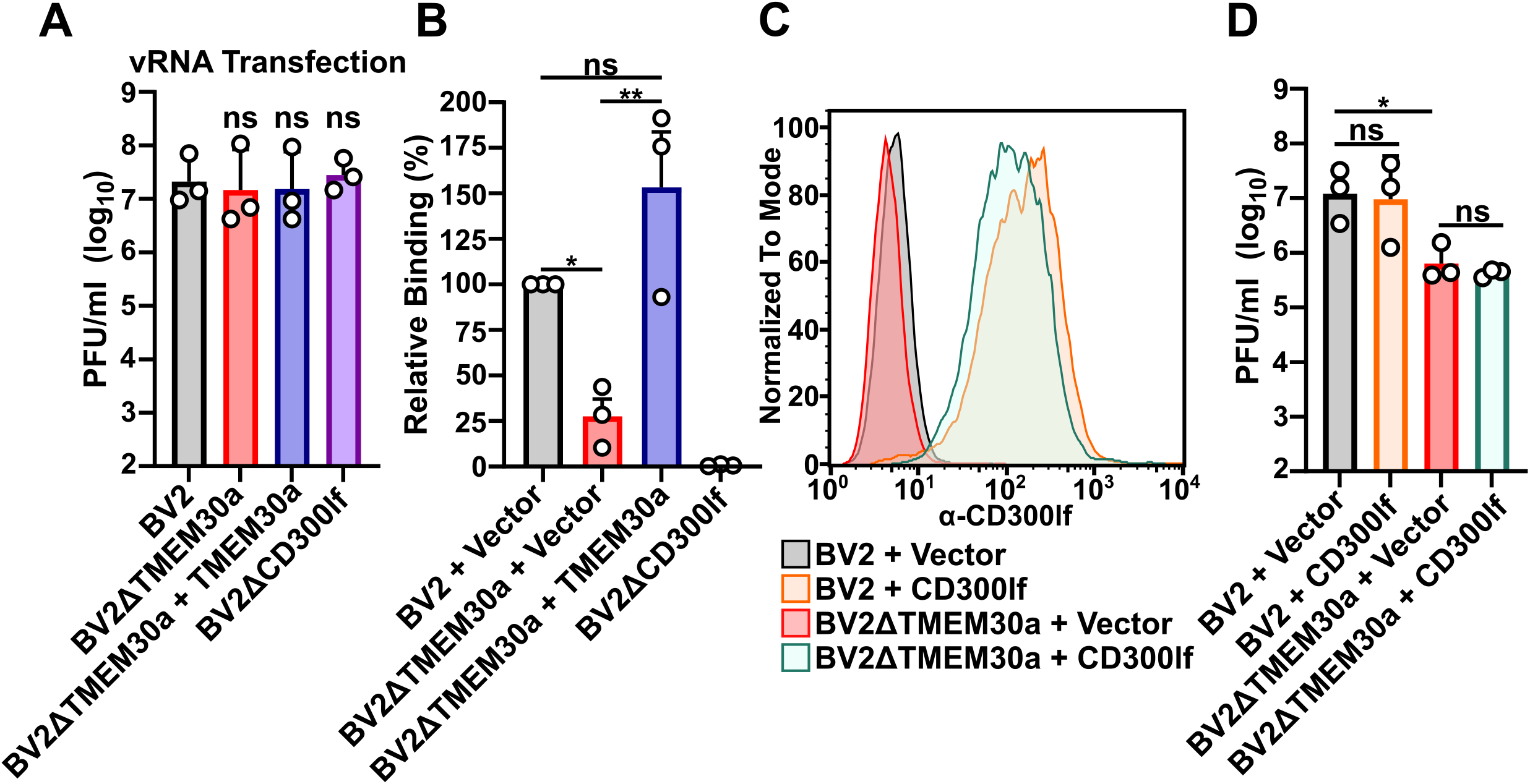
TMEM30a is an MNV entry factor that facilitates viral binding to cells. **A)** Indicated cell lines were transfected with viral RNA from MNV^CR6^ and harvested 12 hours post-transfected. Viral titers were enumerated through plaque assay. **B)** Indicated cell lines were assayed for MNV^CR6^ binding using quantitative polymerase chain reaction and normalized to the mean of BV2 + vector for each experiment. **C)** A representative histogram of surface CD300lf staining of indicated cell lines. **D)** BV2 or BV2ΔTMEM30a cells expressing either an empty vector or CD300lf were challenged with MNV^CR6^ at an MOI of 0.05 and after 12 hours post-infection viral production was enumerated via plaque assay. All data are shown as mean ± S.D. from three independent experiments and analyzed by one-way ANOVA with Tukey’s multiple comparison test. Statistical significance is annotated as follows: ns not significant, * P < 0.05, ** P< 0.01

### Membrane order rather than exoplasmic PS is critical for MNV infection

We next sought to understand how TMEM30a promotes MNV entry. We profiled the exoplasmic PS and PE levels in BV2ΔTMEM30a cells expressing previously reported loss of function mutations to uncouple TMEM30a lipid flipping activities [30]. Fortuitously, we identified cell lines that can be categorized into three distinct plasma membrane states: 1.) cells with no exoplasmic PS or PE, 2.) cells with both exoplasmic PS and PE, 3.) cells with only exoplasmic PE but not exoplasmic PS. More specifically, complementation of BV2ΔTMEM30a cells with TMEM30a reduced exoplasmic PS and PE levels to that of wild-type cells (**Figure 3A-3C**). In contrast, expression of TMEM30a^K308E^ in BV2ΔTMEM30a cells failed to reduce exoplasmic PS and PE levels (**Figure 3A-3C**). TMEM30a^K308E^ is unable to chaperone P4-ATPases out of the endoplasmic reticulum [30]. Lastly, expression of the hypomorphic construct TMEM30a^W260A^ led to a discordance in exoplasmic PS and PE levels in BV2ΔTMEM30A cells [30]. The TMEM30A^W260A^ expressing cells had similar annexin staining levels as wild-type cells indicating the cells had little exoplasmic PS (**Figure 3A and 3B**). However, these same cells had elevated levels of exoplasmic PE as indicated by duramycin staining (**Figure 3A-3C**). This contrast between exoplasmic PS and PE provided a unique opportunity to distinguish the lipid flipping activities of TMEM30a. Indeed, challenging these cell lines with MNV^CR6^ indicates that exoplasmic PS is not sufficient to explain why MNV requires lipid asymmetry. In particular, BV2ΔTMEM30a + TMEM30a^W260A^ cells have elevated exoplasmic PE and were unable to support MNV^CR6^ infection despite normal levels of exoplasmic PS (**Figure 3D**). These data suggest that the lipid flippase activity of TMEM30a is critical for promoting MNV replication and that MNV is more sensitive to exoplasmic PE rather than exoplasmic PS. Consistent with this model, masking the exoplasmic PS on BV2ΔTMEM30a cells by preincubating cells with saturating amounts of annexin V had no impact on viral binding to BV2 or BV2ΔTMEM30a cells (**Figure 3E**). Taken together, these data demonstrate that TMEM30a lipid flipping is critical for MNV recognition of cells and that this dependency on lipid asymmetry is independent of exoplasmic PS.

**Figure 3:**
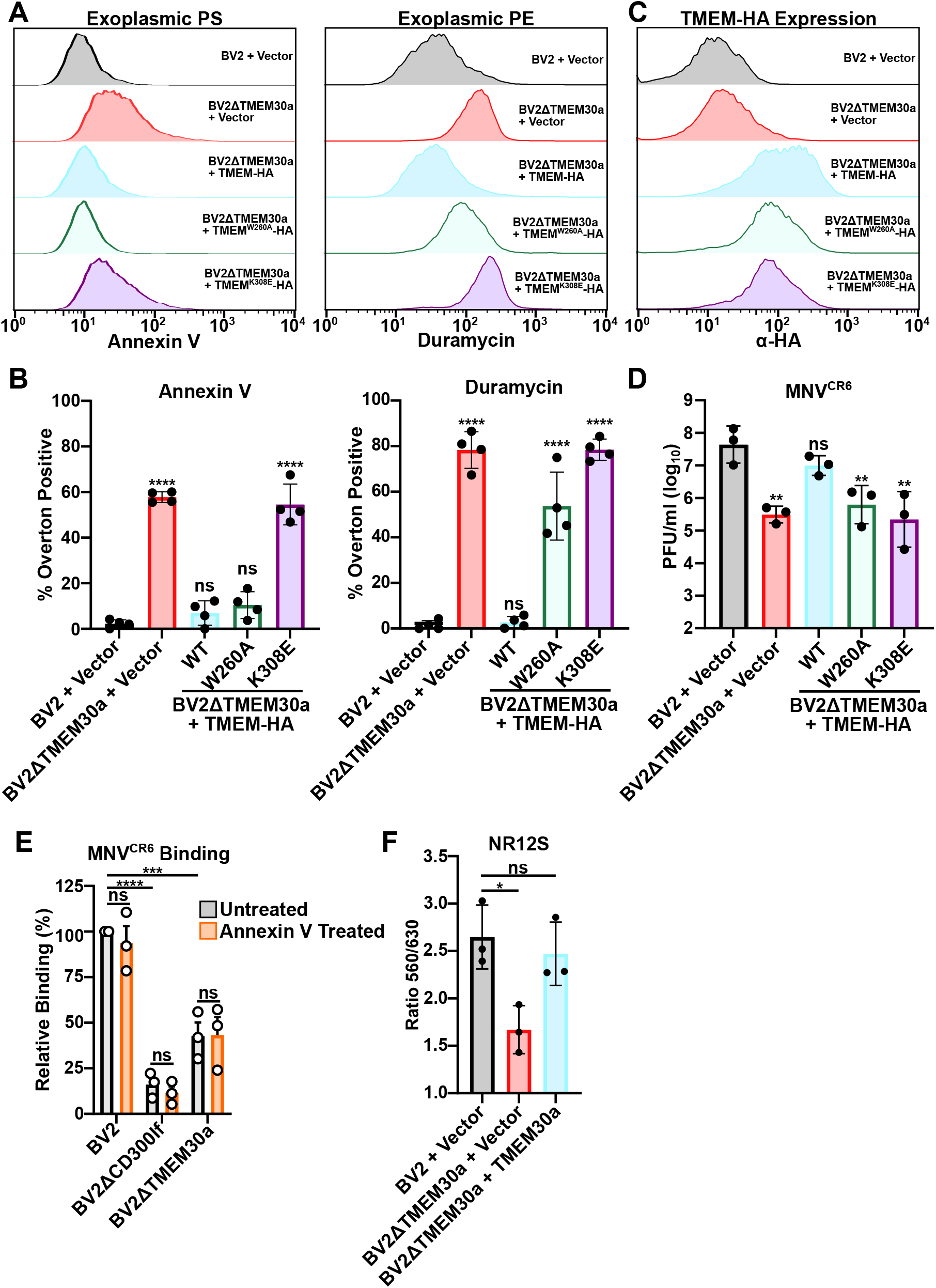
TMEM30a lipid flipping activity of PE but not PS facilitates MNV replication. **A)** Representative histograms from FACS analysis of indicated cell lines stained with Annexin V (left) or Duramycin (right). **B)** Quantification of percent positive cells for Annexin V (Left) or Duramycin (right). Data is from four independent experiments. **C)** Representative histogram of intracellular flow cytometry for TMEM30a-HA of indicated cell lines. **D)** Indicated cell lines were challenged with MNV^CR6^ at an MOI of 0.05 and viral titers enumerated 12 hours post-infection via plaque assay. Data is from three independent experiments. **E)** Indicated cell lines were either incubated either Annexin V or buffer alone on ice prior to performing a binding assay with MNV^CR6^. Within an experiment binding was normalized to BV2 mock treated. Data is from three independent experiments. **F)** Ratio of the emission at 530 nm over 630 nm of indicated cell lines after incubated with NR12S. Data is from three independent experiments. All data are shown as mean ± S.D. from at least three independent experiments and analyzed by one-way ANOVA with Tukey’s multiple comparison test. Statistical significance is annotated as follows: ns not significant, * P < 0.05, ** P< 0.01, *** P< 0.001, **** P< 0.0001

Given that exoplasmic PS was not sufficient to explain MNV infection dependence on TMEM30a, we next explored whether a general disruption of lipid packing and order could be responsible for the phenotype. In addition to headgroup asymmetry, lipids are asymmetrical in lipid saturation which leads to distinct changes in membrane order and fluidity [31]. To characterize the physical properties of membranes we leveraged the outer leaflet selective probe NR12S. NR12S is a derivative of Nile Red whose fluorescence emission shifts towards shorter wavelengths (530 nm) when incorporated into a liquid ordered phase compared to a liquid disordered phase (630 nm) [32]. NR12S fluorescence emission profiles of BV2ΔTMEM30a cells demonstrated a significant increase in membrane disorder as measured by the emission ratio 530nm / 630nm (**Figure 3F**).

Importantly, this phenotype could be restored via complementation with a TMEM30a cDNA (**Figure 3F**). These data indicate that in addition to changes in lipid head group, TMEM30a is required to maintain exoplasmic lipid order, which has the potential to impact MNV binding and entry independent of lipid headgroup.

## Discussion

Here we uncover an unappreciated connection between lipid asymmetry and norovirus entry. Unlike enveloped viruses that disrupt lipid asymmetry to promote their uptake, MNV requires lipid asymmetry to enter cells [5]. These findings add to the growing body of literature that lipids are important mediators of norovirus entry. Sphingolipid biosynthesis maintains CD300lf in a functional confirmation for MNV entry [33]. However, CD300lf staining was not affected by disruptions in lipid asymmetry, pointing to a distinct role for the ordered distribution of lipids in promoting MNV infection (**Figure 2C**). For human norovirus, the addition of ceramide to cells promotes norovirus replication and cellular acid sphingomyelinase promotes a membrane wound repair pathway that enables entry of pandemic human norovirus strains [34, 35]. Whether these features are dependent upon TMEM30a or if human norovirus infection requires TMEM30a warrants investigation.

Noroviruses can be shed long after symptom resolution in asymptomatic individuals [9, 10]. In the MNV system, only a small number of tuft cells in the intestinal epithelium are infected despite high titers of virus being shed in the feces of the mice [23, 36]. Disruption of lipid asymmetry in intestinal epithelial cells dramatically reduces viral persistence in vivo (**Figure 1H and 1I**). Given that TMEM30a deficient cells can support low levels of viral replication in vitro, these data suggests that only a small perturbation to persistent viral infection may be necessary to halt transmission cycles. One limitation of our study is that we are unable to ascertain whether defective MNV entry is the reason why *TMEM30a*^*fl/fl*^*-Vilcre* mice do not support persistent infection of MNV^CR6^ or whether other complex non-cell autonomous factors are responsible. Nevertheless, TMEM30a is required for persistent MNV infection and represents a potential therapeutic target for persistent norovirus infections.

Lipid asymmetry involves the segregation of different lipid classes to distinct leaflets of the lipid bilayer. Disruption of lipid asymmetry in platelets leads to a ratio of two phosphatidylserines for every five phosphatidylethanolamines being exposed [37]. Despite the abundance of exoplasmic PE, most research efforts have focused on the distribution of PS given the number of scavenger receptors that bind PS to facilitate the clearance of dead cells [38]. Exoplasmic PE has been shown to have a similar role for PS or synergize with PS engagement of proteins [39, 40]. In contrast, our work identifies an independent role for exoplasmic PE. Using a TMEM30a mutant that has a partial loss of function (TMEM30A^W260A^), we generated cells that had exoplasmic PE but normal PS distribution (**Figure 3A and 3B**). In this setting, MNV was unable to replicate, suggesting a critical role of PE but not PS in regulating MNV entry (**Figure 3D**). How PE directly or indirectly impacts viral binding to cells is an outstanding question. We propose a model in which changes in the distribution of lipids alter membrane fluidity which in turn reduces MNV binding. This model is supported by the recent finding that decreasing membrane mobility reduced MNV binding due to the low-affinity, high avidity of MNV engaging CD300lf [41, 42]. Indeed, in our in vitro model, BV2ΔTMEM30a cells had a decrease in lipid order, one component that influences membrane fluidity (**Figure 3F**).

Our work has several limitations. First, the mechanism for how the TMEM30a^W260A^ mutant selectively enables PS flipping but not PE is unknown. This phenotype may be cell-type specific or dependent upon the expression levels we achieved in our study. Second, we are currently unable to disentangle lipid saturation asymmetry and head group asymmetry. It is possible that changes in both lipid packing and lipid headgroups are detrimental to MNV infection. Despite over 50 years of research on membrane biology, it remains unclear why an asymmetric bilayer is advantageous to cells [31, 43]. Our research results presented here provide new tools and a physiological system to address this fundamental aspect of cellular physiology.

## Materials and Methods

### Cell Culture

293T (ATCC) and BV2 cells (Kind gift of Dr. Skip Virgin, Washington University) were cultured in Dulbecco’s Modified Eagle Medium (DMEM) with 5% fetal bovine serum (FBS). 3 µg/mL of puromycin (Thermo Fisher) was added as appropriate. BV2ΔCD300lf cells have been described previously [15]. BV2ΔTMEM30a cells were generated at the Genome Engineering and iPSC center at Washington University School of Medicine. BV2 cells were nucleofected with Cas9 and TMEM30a-specific sgRNa (CGGAGATAAACATTTATTGCNGG). Single cell clones were screened for frameshifts by sequencing the target region with Illumina MiSeq at approximately 500X coverage. All cell lines are tested regularly and verified to be free of mycoplasma contamination.

### Plasmids

cDNA for human TMEM30a was obtained from TransOMIC and subsequently cloned into pCDH-MSCV-T2A-Puro vector (System Biosciences) with and without a C-terminal HA-tag. pCDH-MSCV-CD300lf-Flag-T2A-Puro was described previously [15]. All TMEM mutants were generated through splicing by overlap extension PCR. All DNA constructs were sequence verified.

### MNV Assays

MNV^CW3^ (Gen bank accession no. EF014462.1) and MNV^CR6^ (Gen bank accession no. JQ237823) were generated by transfecting molecular clones into 293T cells and amplifying on BV2 cells as described previously [15]. MNV strains CR3, CR7, WU23, and S99 were kind gifts from Dr. Craig Wilen (Yale School of Medicine) [24]. For MNV growth assays, 5 × 10^4^ BV2 cells were seeded in 96-well plates and the next day inoculated with virus and frozen at -80°C at the indicated time point. MNV plaque and entry bypass assays were done as described previously [15]. For isolation of MNV^CR6^ RNA, RNA was purified from cell-free viral preparations using the Direct-zol RNA Miniprep kit (Zymo Research catalog #R2052) according to manufacturer instructions. Purified RNA was verified to be free of infectious MNV particles via plaque assay. 10 µg of RNA was transfected using Lipofectamine 2000 transfection reagent according to the manufacturer’s protocol. Transfected cells were frozen 12 hours later. All infections were done in triplicate in each of at least three independent experiments. MNV^CR6^ binding assays were performed as previously described [33]. Briefly, MNV^CR6^ binding to BV2 cells was performed for 1 hour at 4°C in 0.5 mL DMEM with 10% FBS. BV2 cells were used at a final concentration of 2.5 × 10^6^ cells/mL and MNV was used at an MOI of 1. Cells were centrifuged 500 *g* for 5 minutes at 4°C to remove unbound virus. Cells were washed four times with 1 mL cold DMEM with 10%FBS. The cell pellet was resuspended in 100 µL PBS, and RNA was extracted using the Direct-zol RNA Miniprep kit according to manufacturer instructions. qPCR was utilized to quantify MNV and Actin transcript abundance. Binding assays were conducted in triplicate in at least three independent experiments and binding was normalized within an experiment to the ratio of MNV/Actin and normalized between experiments by setting the relative binding of MNV to WT BV2 cells at 100% as done previously [33].

### Flow Cytometry

For fluorescence-activated cell sorting (FACS) analysis, cells were isolated and probed for either Annexin-V (Biolegend #64095) or Duramycin-LC-Biotin (polysciences catalog#25690-100) in annexin staining buffer (0.01 M Hepes Buffer, 0.14 M NaCl, 2.5 mM CaCl_2_) or PBS respectively. Samples were incubated at 4°C for 30 minutes prior to washing and staining with FITC- or PE-conjugated Streptavidin. The fluorescent intercalator 7-Aminoactinomycin D (7-AAD) (Fisher Scientific catalog#A1310) was used as a control to observe cell-viability. For CD300lf expression, cells were stained with the PE-TX70 antibody (Biolegend #132704) as described previously[33]. For intracellular FACS, cells were fixed and permeabilized with Cytofix/Cytoperm (BD Biosciences) prior to staining Rabbit monoclonal anti-HA antibody (CellSignaling Technologies #3724). At least 20,000 events were collected per condition. Each experiment was performed in triplicate in each of three independent experiments. Cells were analyzed on a FACS Calibur flow cytometer and data analyzed using FlowJo v10 (FlowJo LLC, Ashland, OR).

### Mouse Infections

All mouse experiments were conducted at University of Texas Southwestern Medical Center and approved by the University of Texas Southwestern Medical Center’s Institutional Animal Care and Use committees. We thank the Wellcome Trust Sanger Institute Mouse Genetics Project (Sanger MGP) and its funders for providing the mutant mouse line *Tmem30atm1a(KOMP)Wtsi* and INFRAFRONTIER/EMMA (www.infrafrontier.eu). Funding information may be found at www.sanger.ac.uk/mouseportal and associated primary phenotypic information at www.mousephenotype.org. *Tmem30atm1a(KOMP)Wtsi* mice were first bred to *B6*.*Cg-Tg(Pgk1-flpo)10Sykr/J* (Jax stock #011065) to remove the genetrap cassette and create a conditional knockout mouse. We then bred *TMEM30a*^*fl/fl*^ mice to *B6 Vil-cre* (Jax stock #004586) in a pathogen-free barrier facility including devoid of murine norovirus. 1 × 10^6^ PFU of MNV^CR6^ diluted in 25 µl of DMEM was inoculated per orally to 6-8 week old mice. Immediately after inoculation, mice were singly housed. Fecal pellets were collected at 0, 3, 7, 14, and 21 days post-infection. Mice were euthanized at 21 days post-infection and tissues were harvested and prepared for RNA isolation.

### Quantitative PCR

MNV genome copies in fecal pellets and tissues were determined as previously described. Briefly, RNA was isolated from infected tissues using TRI Reagent (Sigma-Aldrich # T9424) with a Direct-zol kit (Zymo Research) following the manufacturers’ protocols. 1 µg of RNA was used for cDNA synthesis using a High-Capacity cDNA Reverse Transcription kit, following the manufacturer’s protocols (Thermo Fisher Scientific # 4368813). TaqMan quantitative PCR (qPCR) for MNV was performed in triplicate on each sample and standard with forward primer 5’-GTGCGCAACACAGAGAAACG-3’, reverse primer 5’-CGGGCTGAGCTTCCTGC-3’, and probe 59-6FAM-CTAGTGTCTCCTTTGGAGCACCTA-BHQ1-3’. TaqMan qPCR for Actin was performed in triplicate on each sample and standard with forward primer 5’-GATTACTGCTCTGGCTCCTAG-3’, reverse primer 5’-GACTCATCGTACTCCTGCTTG-3’, and probe 5’-6FAM-CTGGCCTCACTGTCCACCTTCC-6TAMSp-3’. RNA from infected fecal pellets was isolated using RNeasy Mini QIAcube Kit (Qiagen # 74116) and cDNA synthesis was performed using M-MLV Reverse Transcriptase kit (Invitrogen # 28025013) using manufacturer’s protocols.

### NR12S

5 × 10^4^ BV2 cells were seeded in black walled 96-well plates and the following day media was removed and 100 nM NR12S (Tocris) diluted in DMEM without FBS was added to cells at room temperature for 20 minutes. Media was removed, and PBS was added to cells. Samples were read on a BioTek Cytation5 plate reader with an excitation of 520 nm and separate emission readings at 560 nm and 630 nm.

## Acknowledgements

We would like to thank Craig Wilen, Chaitanya Dende, Neal Alto, and Arun Radhakrishnan and all members of the Orchard Lab for helpful discussions. B.M.S. was supported by the UT Southwestern Molecular Integrative Immunology Training Grant (T32AI005284). R.C.O. was supported by NIH grant R35GM142684.

## Author Contributions

B.M.S., L.R.P., and M.C.O. designed the project, performed experiments and helped draft the paper. R.C.O. conceptualized the project, provided supervision, and helped write the paper. All authors read and edited the manuscript.

